# Modelling optically pumped magnetometer interference as a mean (magnetic) field

**DOI:** 10.1101/2020.11.25.397778

**Authors:** Tim M Tierney, Nicholas Alexander, Stephanie Mellor, Niall Holmes, Robert Seymour, George C O’Neill, Eleanor A Maguire, Gareth R Barnes

## Abstract

Here we propose that much of the magnetic interference observed when using optically pumped magnetometers can be modeled spatially as a mean (magnetic) field. We show that this approximation reduces sensor level variability and substantially improves statistical power. This model does not require knowledge of the underlying neuroanatomy nor the sensor positions. It only needs information about the sensor orientation. Due to the model’s low rank there is little risk of removing substantial neural signal. However, we provide a framework to assess this risk for any sensor number, design or subject neuroanatomy. We find that the risk of unintentionally removing neural signal is reduced when multi-axis recordings are performed. We validated the method using a binaural auditory evoked response paradigm and demonstrated that the mean field correction increases reconstructed SNR in relevant brain regions in both the spatial and temporal domain. Considering the model’s simplicity and efficacy, we suggest that this mean field correction can be a powerful preprocessing step for arrays of optically pumped magnetometers.

## 1. Introduction

As Optically Pumped Magnetometers (OPMs) become more sensitive (Kominis, Kornack, Allred, & Romalis, 2003), there is an increasing need to develop methods that minimize the magnetic interference they experience. A number of distinct approaches to this problem have been developed.

Hardware developments to mitigate interference include active shielding systems to null low-frequency fields around the participant’s head (Holmes et al., 2019; Iivanainen, Zetter, Grön, Hakkarainen, & Parkkonen, 2019). This approach is attractive as it allows simplification of the helmet design (to comprise just the magnetometers). Furthermore, these coils have made feasible the imaging of neural responses during participant movement (Boto et al., 2019; Holmes et al., 2018, 2019). Whilst these coils prevent artefacts in the region of 0–1 Hz that would move the sensors outside their dynamic range, they do not correct for the subtler modulations of interference encountered when someone rotates their head. This modulation of interference could theoretically be corrected with high dynamic range field nulling coils but this further complicates the hardware development.

Another hardware option is to configure the sensors as gradiometers which have excellent noise suppression (Colombo et al., 2016; Nardelli, Perry, Krzyzewski, & Knappe, 2020; Sheng et al., 2017) and are capable of unshielded neural recordings (Limes et al., 2020). In contrast, atomic magnetometers (Osborne, Orton, Alem, & Shah, 2018) typically require considerable active and passive shielding for successful MEG recordings (Barry et al., 2019; Lin et al., 2019; Tierney et al., 2018, 2020). However, the appeal of magnetometers lie in their simplicity of construction, compact size and extreme sensitivity (Allred, Lyman, Kornack, & Romalis, 2002; Kominis et al., 2003). Furthermore, gradiometers necessarily must have a baseline that is 1-2 times larger than the distance to the brain region of interest (Hämäläinen, Hari, Ilmoniemi, Knuutila, & Lounasmaa, 1993). This constraint may place a limit on the wearability of atomic gradiometer systems, and consequently compact magnetometer designs for wearable systems are more attractive.

In addition to hardware developments, there are a number of software approaches that can be used to reduce interference. Perhaps some of the most widely used methods are the Signal Source Separation (SSS) method (Taulu & Kajola, 2005) and the Dual Signal Subspace Projection (DSSP) method (Sekihara et al., 2016). Both methods aim to partition the data into separate subspaces that originate from inside the brain and outside the brain. Once these subspaces are defined the temporal intersection of these subspaces can be used to further reduce interference (Golub & Van Loan, 1996). When this temporal extension is used with SSS it is referred to as tSSS (Taulu & Hari, 2009). The methods diverge in the form of the basis sets used to represent the neural and external subspace. SSS uses a set of spherical harmonics and considers the low (spatial) frequency terms to originate from outside the head. DSSP uses the eigenmodes of the lead fields to define the neural space while the nullspace of the neural space defines the external space. Both methods have proved very useful in dealing with challenging interference such as that from vagus nerve stimulators or deep brain stimulators (Cai et al., 2019; Kandemir, Litvak, & Florin, 2020).

However, these methods make a number of assumptions that should be considered before use with small channel (< 50) OPM systems. For instance, inherent to DSSP is the assumption that the rank of the data is much greater than the rank of the lead fields. This is clearly the case in cryogenic MEG systems that may have 300 sensors and a lead field rank in the range of 50-100 (Iivanainen et al., 2020; Nenonen, Taulu, Kajola, & Ahonen, 2007; Tierney, Mellor, et al., 2019). In OPM systems this may not be the case and the rank of the lead fields may be comparable to the rank of the data because typical systems currently operate with fewer than 50 sensors (Hill et al., 2020).

SSS may suffer from an opposing issue in OPM systems, namely that the rank of the external subspace (typically 16) may share significant variance with the leadfields of small channel systems (< 50 sensors). To mitigate this issue, one could use lower order spherical harmonics as the basis set defining the external subspace. Intriguingly, the lowest order spherical harmonic (the constant term) is absent from the SSS basis set on the grounds that it reflects the field from monopoles (Chella, Zappasodi, Marzetti, Penna, & Pizzella, 2012; Nurminen, Taulu, & Okada, 2008; Taulu & Kajola, 2005).

While we do not expect any interfering magnetic field to be exactly constant in space, we may approximate this interfering field as a spatially constant field. Such an approximation may not be physically correct, but we will show it to be useful for interference mitigation for two reasons. Firstly, the model is very low order (rank 3 –the orthogonal spatial axes) and should therefore share minimal variance with the lead fields. Second, because of its simplicity, it can be updated in real time and track interference that is modulated by participant movement. We will motivate its use in the next section with a toy problem of parallel sensors.

## 2. Theory

### 2.1 Magnetic interference is approximately spatially constant

This effect is the basis of all gradiometer based MEG systems and has been extensively described elsewhere (Hämäläinen et al., 1993; Vrba & Robinson, 2002). We restate it here for completeness. The interfering signal measured by a sensor (*S*_1_) on the head is inversely proportional to the square of the distance (*r*) to the interfering field’s source.

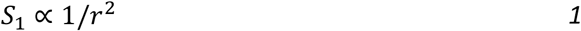

The relative signal (*S*_2_) experienced by a second (parallel) sensor displaced a distance *r* + *h* from the interference source can be described as

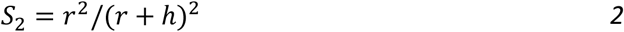

By way of example, both *S*_1_ and *S*_2_ could be sensors either side of a participant’s head. In this case *h* ≈ 20 *cm*. When the distance to the interference is much greater than the head size (*r* >> *h*), the relative signal is approximately constant as a function of space

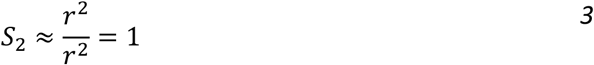

As such, we argue that much of the interference encountered in OPM systems can be described by a spatially constant term (a mean field). It should be noted that this term will automatically be removed in gradiometer systems and will be of most benefit to systems based on magnetometers.

Although this basis set is technically not physically correct (magnetic interference is not constant across space), it will be shown to be a useful approximation. This spatially constant term can be encoded in a *n* × 3 matrix (*N*) that is simply the row-wise concatenation of the *n* unit normals representing the sensors’ sensitive axes 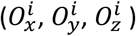. The orientation vectors encode the mean field because these vectors predict the proportion of a constant field experienced by a sensor. The superscript *i* is indicative of the sensor index.

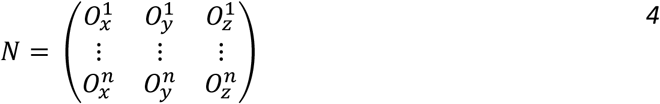

This term should explain the majority of interference distant sources when the participant is stationary. We note that when a participant moves, one could incorporate linear gradient terms. These would be quite beneficial for very low frequency studies (<3Hz), designing a model informed closed loop OPM systems (Nardelli et al., 2020) or for informing coil currents in active shielding systems (Holmes et al., 2019). However, for simplicity we will ignore these higher order terms here and just focus on the benefits of using a very simple mean field approximation.

### 2.2 The separation of signal and interference

Once the basis set (*N*) is defined the *n* × *n* mean field projection matrix (*M*) can be constructed as follows

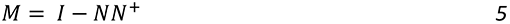

Where *I* signifies the identity matrix and *N*^+^ signifies the pseudoinverse of *N*. This matrix (*M*) projects the *n* × *t* (number of time points) data matrix (*Y*) on to the nullspace of the mean field which will approximately equal the neural subspace (*S_neural_*) for distant interference sources.

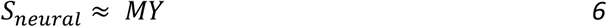

This approach is a simple linear regression (of *N* on to *Y*) and can be viewed as a case of Signal Space Projection - SSP (Uusitalo & Ilmoniemi, 1997) for non-orthonormal basis sets. It should be noted that this approach performs an independent fit of the interference at every time point. As such the model adapts to the temporal non–stationarities in interference.

## 3. Methods

### 3.1 Empty room noise demonstration

We postulated that much of the interference magnetometers experience can be modelled as a mean field. All measurements were made in the new UCL magnetically shielded room specifically designed for OP-MEG. The shielded room, constructed by Magnetic Shields Ltd, has internal dimensions of 4377mm x 3377mm x 2182mm and is constructed from two inner layers of 1mm mumetal, a 6mm Copper layer, and then two external layers of 1.5mm mumetal.

We provide empirical demonstration of this by assessing the ability of the basis set, *N* to mitigate interference in an empty room noise recording. We placed 31 Gen-2 QuSpin OPMs in a participant specific scanner cast (Boto et al., 2016). We recorded 3 minutes of empty room noise data with a 16-bit precision ADC (national instruments) at 6000 Hz sampling rate. We record both the radial and tangential fields from the sensors to double the effective channel count and increase the degrees of freedom. We performed the analysis as described in section 2 using *N* as the basis set.

### 3.2 Lead field errors

Any spatial basis set will share variance (even if only by chance) with the lead fields (the basis set defining the neural subspace). Therefore, a certain percentage of brain signal may be removed following this (or any) regression. To directly assess how much variance is shared between the mean field and the lead fields, we regress the mean field *N*) on to simulated lead fields and calculate the variance explained. The signal loss in decibels (dB) can be calculated as 10 log_10_(1 — *VE*), where *VE* is the fraction of shared variance between the lead fields and *N*.

For the simulation, the sensor array defining the lead fields was a custom made scanner cast (Boto et al., 2016) with 72 sensor slots. For every sensor position two sensor orientations were simulated (one radial and one tangential to the head). The result was 72 sensors and 144 channels. The brain mesh used to generate these leadfields was the MNI canonical mesh available in SPM12, warped to the anatomy of the individual the custom scanner cast was made for. The separation between vertices is approximately 5 mm on average. The orientation of the source was defined by the surface normal of the cortical mesh at that location. The forward model was the Nolte single shell model (Nolte, 2003) and sensors were assumed to be point magnetometers.

### 3.3 Sensor level analysis

Here we show how the proposed approximation can improve the statistical power of sensor level analyses, even during participant motion (~45 degrees rotation) using an auditory evoked response paradigm. One male, aged 26 years, participated in this study and gave informed written consent in line with UCL ethics. The auditory tones had a duration of 70ms (5ms rise and fall times) and frequencies of 500-800Hz in steps of 50Hz. The inter-stimulus interval was 0.5s. Stimuli were presented via Psychopy (Peirce 2009), through MEG-compatible ear tubes with etymotic transducers, and the volume was adjusted to a comfortable level, as specified by the participant. A total of around 1400 individual auditory tones were presented. During the experiment, the participant was instructed to continually, slowly rotate their head by 45 degrees in any direction that was comfortable and to ignore the auditory tones. This was done to deliberately create rotation induced non-stationarities in the recorded data. No motion tracking was performed.

Data were acquired with the same 31 sensors (62 channels) in section 3.1 at 6000Hz using a national instrument 9205 ADC (16 bit system) and subsequently downsampled to 600Hz. The same mean field correction was applied to the data as in section 3.1. Data were then band passed filtered between 2 and 40Hz with a notch at 50Hz. The data were averaged across trials to observe an evoked response. A one sample student t-test was conducted at each time point across trials. Data were corrected for multiple comparisons using Bonferroni correction. For comparison, the same analysis pipeline was repeated but without the use of the mean field correction.

### 3.4 Source level analysis

To examine the effect of the mean field correction at the source level, we reconstructed the source space time courses using Minimum Norm (Hämäläinen et al., 1993) as implemented in SPM12 (Friston et al., 2008; Lopez, Litvak, Espinosa, Friston, & Barnes, 2014) for both the mean field corrected data and the uncorrected data.

We then (in a similar manner to the sensor level) constructed one-sample t-tests but this time focusing on the M100 evoked response. The resulting statistical parametric map was smoothed with a 20mm Gaussian kernel and corrected for multiple comparisons using Bonferroni correction. The squared t-statistic (across trials) represents the SNR (power) of the evoked response and can be interpreted as an F-statistic showing at what time points SNR is greater than 0. We calculate these SNR-time series for both the mean field corrected and uncorrected data at the region of highest statistical power (global maximum). We also calculate how this SNR varies with distance from the global maximum to measure the Full Width at Half Maximum (FWHM) of the SNR.

### 3.5 Software

All analysis was carried out using the SPM12 (https://www.fil.ion.ucl.ac.uk/spm/) software package and custom Matlab scripts. All software is available on request from the corresponding author and will be made freely available via GitHub (https://github.com/tierneytim/OPM).

## 4. Results

### 4.1 Empty room demonstration

In the empty room recordings –see Figure 1-the drift (<1Hz) and 50 Hz components are reduced by a factor of 10, whereas the vibration components within the 3 and 20 Hz band are reduced by up to a factor of 5 (15dB). It is important to note that this improvement was achieved by using only the 3 regressors of the basis *N*. The noise floor reached 10fT at higher frequencies. These reductions provide an empirical verification that the mean field approximations does indeed explain a majority of variance in the magnetic interference encountered in our magnetically shielded room.

**Figure 1.**
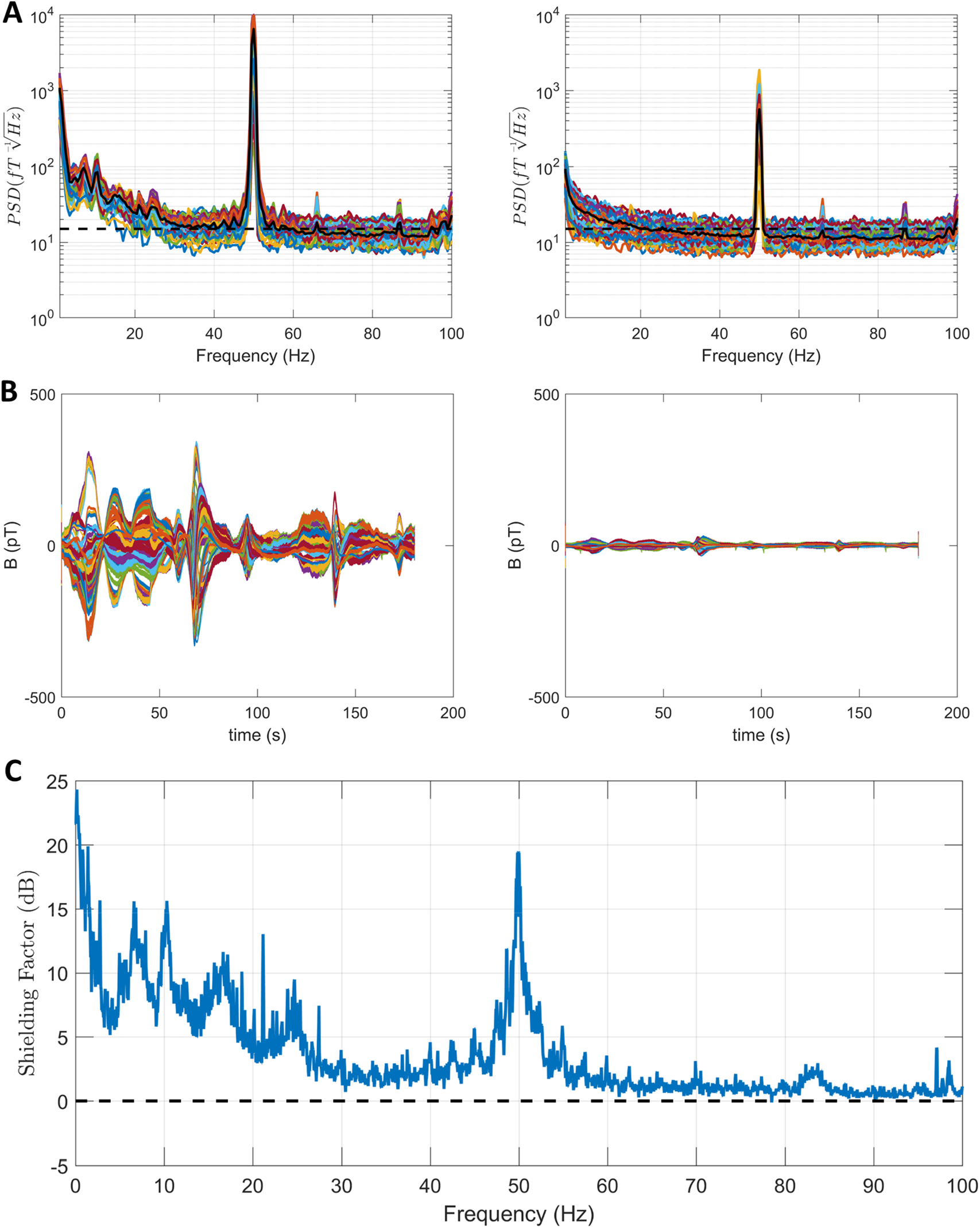
Mean field approximation for empty room noise data. In (A) the power spectral density (rms) is displayed for both uncorrected data (left) and corrected data (right). The average power spectral density is highlighted in black. The dotted black line is at 15 fT. Large sources of interference (drift, 50 Hz) experienced a 10-fold reduction in magnitude while nearly 5-fold reduction in interference was observed between 5 and 20 Hz. In (B) representative time segments are shown. The uncorrected data in the left panel show that the environmental noise within the room could change quickly by a few hundred picotestla. Whereas in the corrected time series (right) these changes were reduced to the order of tens of picotesla. In (C) the average shielding factor across the 62 channels in decibels (dB) is plotted as a function of frequency. There is a positive effect across the entire bandwidth analysed (0-100 Hz).

### 4.2 Lead field errors

We next examined whether the proposed mean field correction attenuated useful signal as well as interference. As is apparent in Figure 2, the signal loss was, on average, less than 0.5dB for an array consisting of both radial and tangential OPMs. It reaches a maximum of ~1.5dB for the combined array and, as such, the reduction in noise observed in Figure 1 outweighs any signal loss. What is perhaps most interesting here is that the signal loss is much lower for a dual-axis recording as opposed to recordings from either axis individually. There is also a clear pattern of increased signal loss at depth with single axis recordings, although, this effect is mitigated by dual-axis recordings.

**Figure 2.**
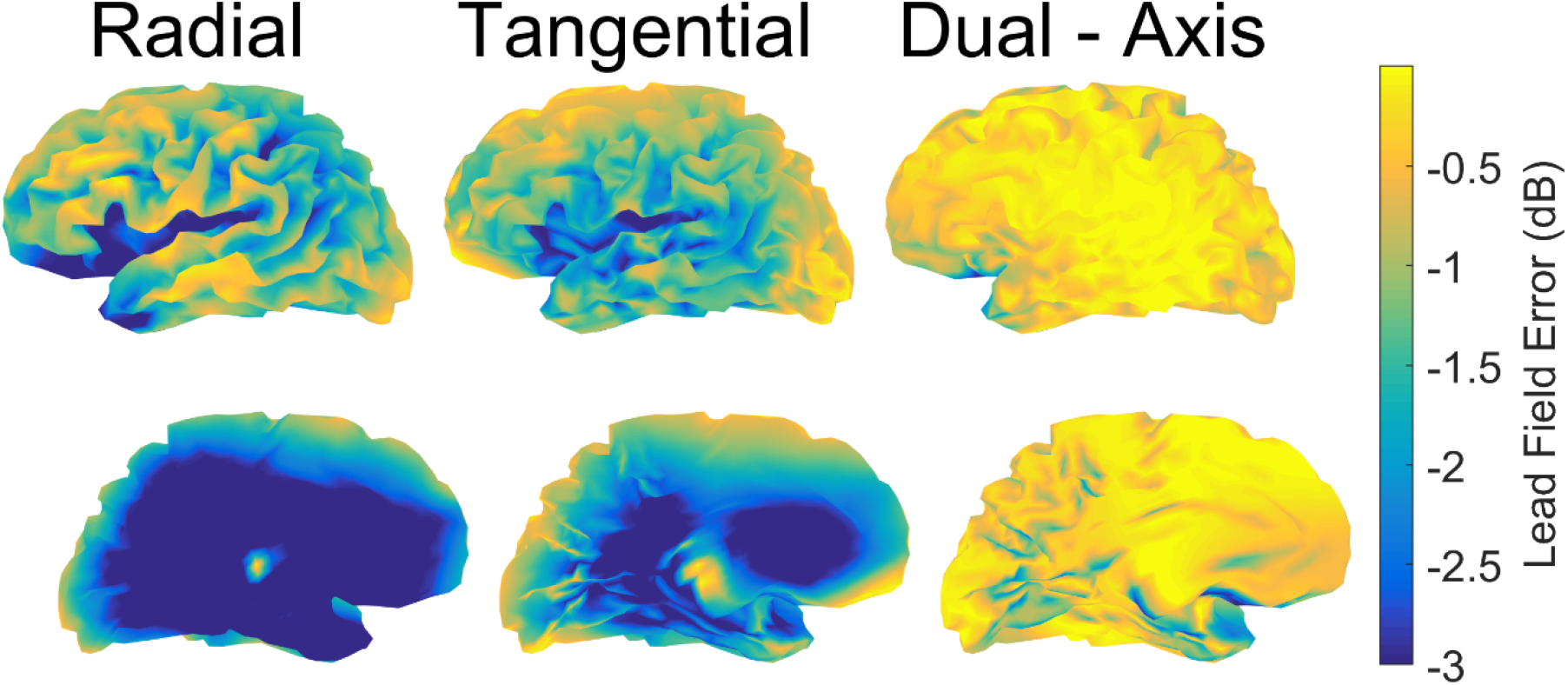
Maximal lead field error due to mean field approximation. For the 72 sensor system the lead-field error (expressed as signal loss in dB, more negative values indicating greater signal loss) is shown for sensors measuring radially, tangentially and in dual axis mode. It is clear that the deeper sources share more variance with the mean field basis set but this effect is greatly diminished when data are collected in dual axis mode.

### 4.3 Sensor level analysis

As shown in Figure 3, we assessed OPM measured auditory evoked responses during movement (~45 degrees rotation). The sensor level evoked response (in Figure 3A and 3B) was obscured across many sensors due to trial to trial variation. This is reflected in the t-statistics which either did not achieve significance or were weakly significant for the early 100ms component. However, when the mean field correction was applied, the evoked response became much clearer (Figure 3C). This reduction in variation was reflected in the increased magnitude of the t-statistics (Figure 3D).

**Figure 3.**
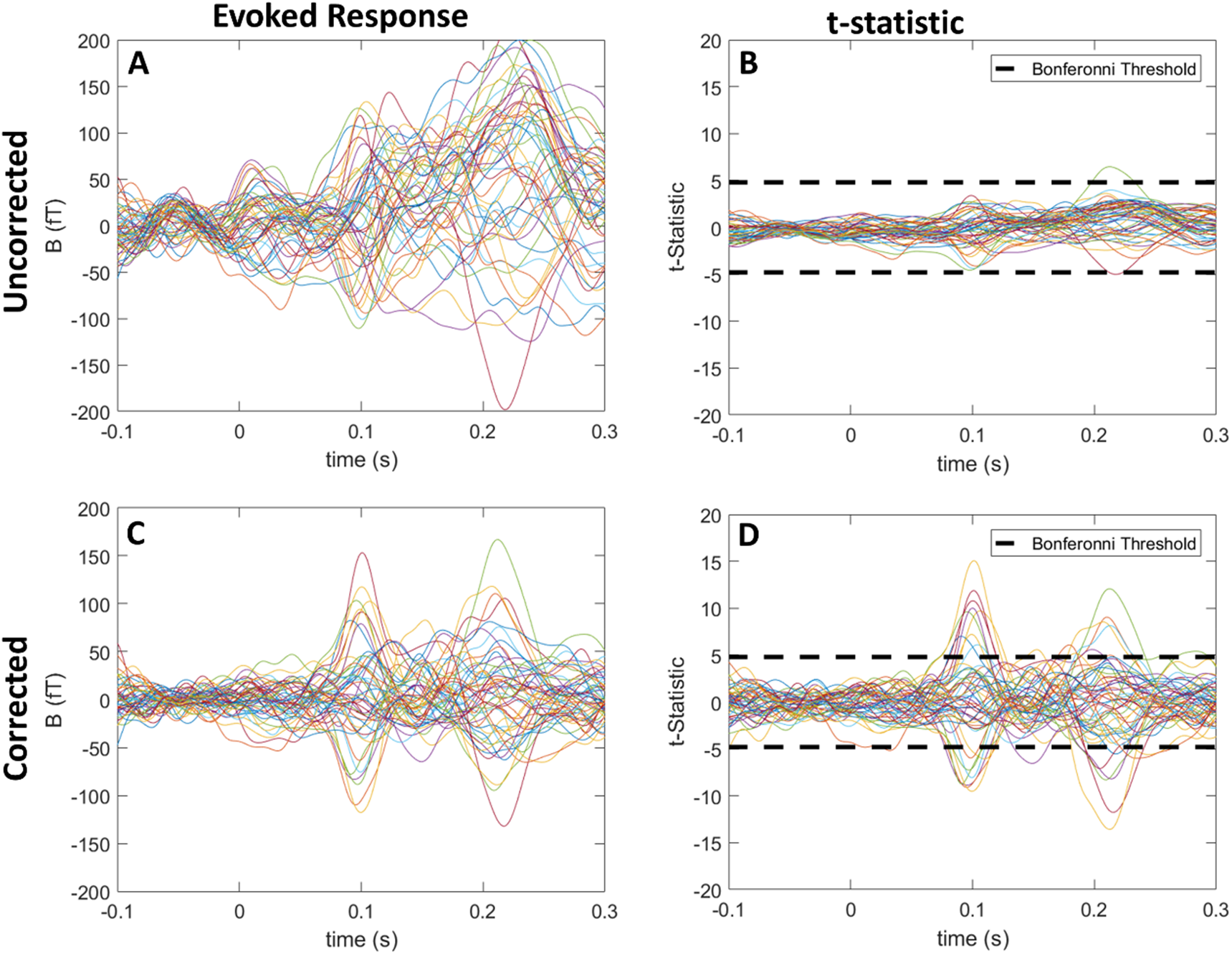
Interference correction in the presence of motion. In (A) and (B) the evoked response and associated t-statistics are shown for the auditory evoked response when no correction is applied. The 100ms and 200ms response are difficult to discern from the data and the statistical efficiency is poor due to the high level of variation. However in (C) and (D) when the mean field approximation is applied both the 100ms and 200ms response are clearly visible in both the evoked response and the t-statistic.

### 4.4 Source level demonstration

The results were encouraging when we examined the source level. The SPMs were both broadly similar for corrected and uncorrected data but the mean field corrected data had more suprathreshold vertices and higher statistical power (Figure 4A and 4B). When we looked directly at how the SNR changed with space (Figure 4C) we observed that the FWHM of the SNR was comparable between both methods, but the SNR was higher for the mean field corrected data. We also directly calculated the SNR across trials (power) at every time point for both methods. This can be interpreted as an F-statistic and clearly showed that the mean field corrected data had better source level SNR than the uncorrected data (Figure 4D) at the time points of high signal (100ms and 200ms).

**Figure 4.**
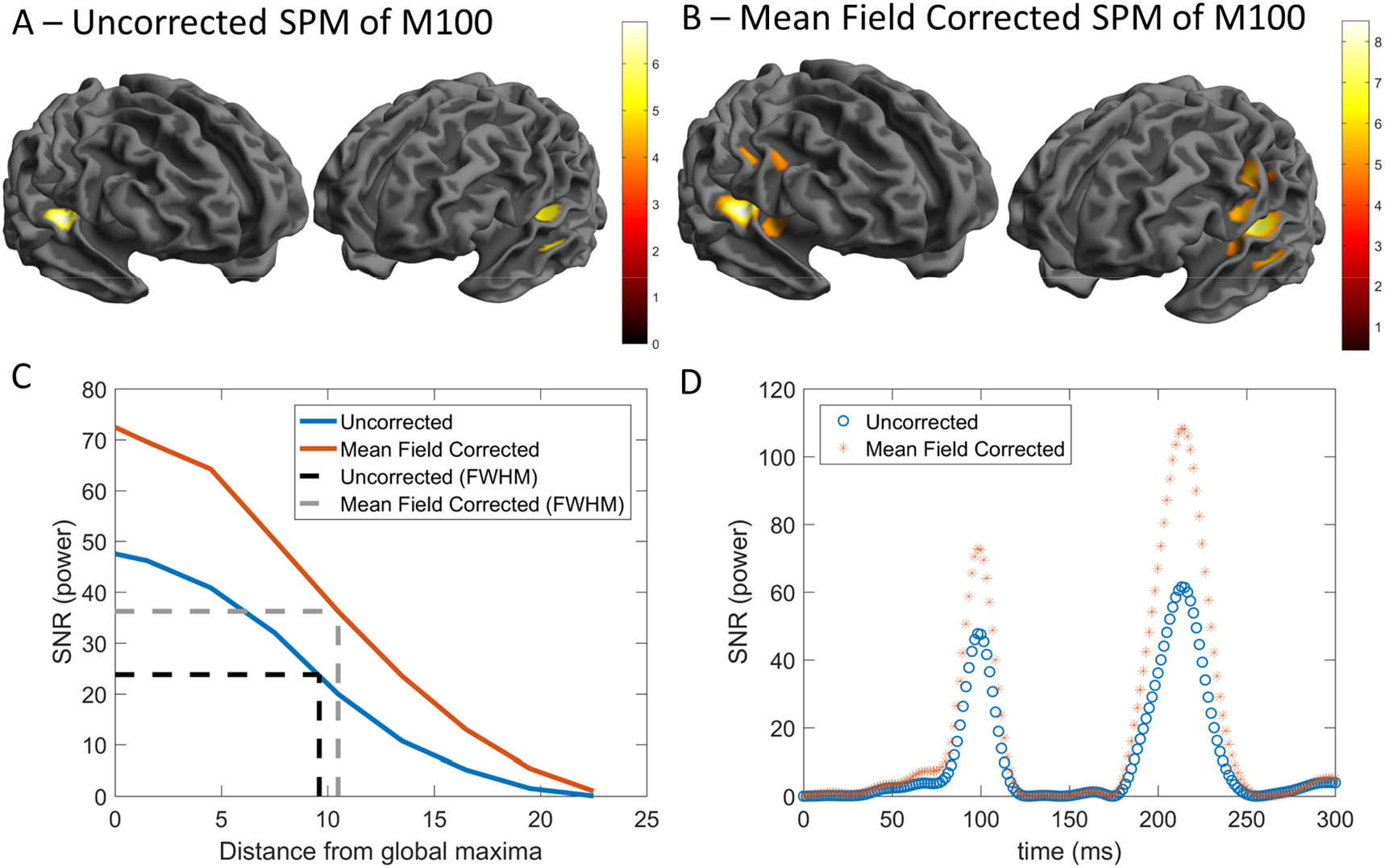
Source level results. In (A) and (B), statistical parametric maps (t-statistic) are shown for the 100ms response to auditory stimuli for both the uncorrected and mean field corrected data respectively. Both situations resulted in bilateral maxima observed in auditory cortex. In (C) the SNR as a function of space is shown. Both methods had comparable FWHMs but the mean field corrected data had better SNR. In (D) the SNR (power) is shown over time for both methods. The mean field method reconstructed more power in the auditory cortex with higher SNR for both the 100ms component as well as the 200ms component.

## 5. Discussion

In this study we demonstrated that a mean field correction provides a simple but powerful approach for the reduction of interference observed by OPMs in both the temporal and spatial domains.

The correction we propose adds to the existing model-based software approaches for the separation of signal from interference. The attraction of using such a low order model lies in both the simplicity of implementation and the low likelihood of removing neural signal. As sensor numbers in OPM arrays are typically much lower than cryogenic MEG systems, the likelihood that any spatial basis set will explain some neural signal by chance is increased. As such, default settings for current spatial desnoising algorithms may not be appropriate, and will need to be adjusted for OPM experiments. Here we provide an explicit framework by which it is possible to calculate the worst-case shared variation for any given array or sensor design and any user specified basis set.

This framework could easily be extended to incorporate a temporally extended mean field correction, similar to tSSS (Taulu & Hari, 2009) but this should be done with some caution. As already noted, for some arrays there will be a non-negligible correlation between a basis set and a lead field. As such, some neural activity will exist in the intersection of these subspaces. This will therefore distort neural activity and should be considered carefully on a case by case basis. The temporal extensions should, arguably, only be used if one can demonstrate that the spatial basis defining the external subspace is nearly orthogonal to the leadfields.

An alternative and powerful model based approach lies in the use of DSSP (Sekihara et al., 2016) which uses the eigenmodes of the lead fields as a spatial basis set to explain the data (effectively modelling the neural subspace at the sensor level). As it is directly modelling the neural subspace it typically rejects less neural data than tSSS (Cai et al., 2019). However, it relies on having an accurate forward model. Neither SSS nor the mean field correction suggested here require a forward model. In fact, the mean field correction only requires knowledge of the relative sensor orientations, whereas SSS requires knowledge of the sensors positions and orientation.

More data driven approaches lie in the use of adaptive source reconstruction techniques such as beamformers (Belardinelli, Ortiz, Barnes, Noppeney, & Preissl, 2012; Van Veen & Buckley, 1988). In principle, for stationary participants, the mean field correction will be of little benefit for beamformer studies (unless the interference covariance changes over time). However, for moving participants, the beamformer will be less effective at removing correlated interference because the estimate of the covariance matrix will be inefficient and biased due to movement induced non-stationarities (although see (Woolrich et al., 2013) for a non-stationary implementation). Ultimately, the mean field correction suggested here offers a compromise between model complexity, variance shared with the leadfields, ease of implementation and non-stationary interference reduction.

While the proposed correction outlined here can help improve the quality of data in OPM experiments using a very low order model, it has some limitations. Most notably, the basis set will share some variation with the leadfields. This shared variation gets smaller with increased number of sensors and simultaneous multi-axis measurements but it nevertheless still exists. The effect of this will depend on the array geometry, sensor design and sensor number. The results here clearly point to the utility of multi-axis measurements to help mitigate this problem. With regards to sensor numbers one can reproduce the analysis of Figure 2 for any channel count/positioning or subject specific anatomy and weigh up the expected signal loss with the observed interference reduction.

An issue associated with array geometry is the requirement for accurate knowledge of the sensors’ sensitive axis. This may be slightly different from the physical orientation of the sensor due to the presence of cross-talk (Tierney, Holmes, et al., 2019) or imperfect on board coil design. However, such issues can be reduced by operating the sensors with coils specifically designed to reduce cross-talk (Nardelli, Krzyzewski, & Knappe, 2019) or by the use of data driven approaches which can learn sensor sensitive axes from the data (Duque-muñoz et al., 2019).

Necessarily, OPMs operate with the aid of on board magnetic coils to maintain zero field at the sensor (Osborne et al., 2018; Tierney, Holmes, et al., 2019). If the magnetic field from these on board coils is not updated as the field at the sensor changes, there will be a component of the motion artefact that is a function of this initial sensor specific magnetic field (e.g. when someone rotates their head the magnetic field designed to keep zero field in one orientation will be incorrectly applied to a different orientation). We do not investigate this effect here but note that it could be mitigated by operating sensors in a closed loop mode (Nardelli et al., 2020), learning this field profile directly from the data itself or by utilizing active shielding so as to keep these values to a minimum (Holmes et al., 2018, 2019; Iivanainen et al., 2019).

While these issues will be the subject of future work, the data presented here clearly show that a simple mean field correction can mitigate much of the interference observed in OPM recordings and improve statistical power both temporally and spatially at sensor and source level. This approximation benefits from multi-axis measurement and, in this case, has minimal negative impact on the neural subspace. These features, coupled with its ease of implementation and lack of reliance on knowledge of the underlying neuroanatomy, render this an appealing and powerful preprocessing step for arrays of OPMs.

## Acknowledgements

This work was supported by a Wellcome collaborative award to GRB, Matthew Brookes and Richard Bowtell (203257/Z/16/Z, 203257/B/16/Z). SM was funded through the EPSRC-funded UCL Centre for Doctoral Training in Medical Imaging (EP/L016478/1). The Wellcome Centre for Human Neuroimaging is supported by core funding from the Wellcome Trust (203147/Z/16/Z). EAM is funded through a Wellcome Principal Research fellowship (210567/Z/18/Z). We also would like to thank the Matthew Brookes, Richard Bowtell, Vishal Shah and David Woolger and their respective teams at University of Nottingham, QuSpin and Magnetic Shields Ltd for their support throughout the development of our OPM system.

